# Convergent representations and spatiotemporal dynamics of speech and language in brain and deep neural networks

**DOI:** 10.1101/2024.12.28.630582

**Authors:** Peili Chen, Shiji Xiang, Linyang He, Edward F. Chang, Yuanning Li

## Abstract

Recent studies have explored the correspondence between single-modality DNN models (speech or text) and specific brain networks for speech and language. The key factors underlying these correlations and their spatiotemporal evolution within the brain language network remain unclear, particularly across different DNN modalities. To address these questions, we analyzed the representation similarity between self-supervised learning (SSL) models for speech (Wav2Vec2) and language (GPT-2), against neural responses to naturalistic speech captured via high-density electrocorticography. Our results indicated high prediction accuracy of both types of SSL models relative to neural activity before and after word onsets. It was the shared components between Wav2Vec2.0 and GPT-2 that explained the majority portion of the SSL-brain similarity. Furthermore, we observed distinct spatiotemporal dynamics: both models showed high encoding accuracy 40 milliseconds before word onset, especially in the mid-superior temporal gyrus (mid-STG), which can be explained by the shared contextual components in the SSL models; the Wav2Vec2.0 also peaked at 200 milliseconds after word onset around the posterior STG, which was mainly attributed to the acoustic-phonetic and static semantic information encoded in the SSL models. These results highlight how contextual and acoustic-phonetic cues encoded in DNNs align with spatiotemporal neural activity patterns, suggesting a significant overlap in how artificial and biological systems process linguistic information.

## Introduction

Since David Marr’s three-levels of analysis^1^, understanding brain processing through computational models stands as a pivotal endeavor in neuroscience, bridging the gap between biological cognition and artificial intelligence^2,3^. These models offer a unique lens to examine complex neural functions, providing insights that are often inaccessible through traditional experimental techniques alone. Computational models facilitate the translation of neurological data into quantifiable patterns that can be analyzed systematically. By mimicking the way the human brain processes information, computational models such as deep neural networks (DNNs) allow researchers to dissect the intricate mechanisms underlying speech, language, and cognitive functions^4–7^. In particular, modern DNN models have allowed researchers to build encoding models that predict neural activity and behavior patterns, shedding light on the computational principles of the cognitive processes^6,8^ .

The past decade has witnessed remarkable breakthrough in artificial intelligence models for speech and language processing. Transformer-based self-supervised models have revolutionized the field of speech and language processing^9–11^. These models achieved state-of-the-art performance in speech and language tasks, such as automatic speech recognition^10,12,13^, dialog, text generation and translation^11^. Furthermore, recent studies have also employed these different models as neural encoding models. Comparing to traditional feature encoding models based on manually defined ad-hoc features derived from theoretical hypotheses^14,15^, these models yield better accuracy in predicting neural activity^16–18^. These studies demonstrate that the internal representations and contextual computations in these DNN models linearly correlated to neural activity in different brain areas underlying speech^16,17^ and language processing^18^ .

Despite having achieved high brain-prediction accuracy, these Transformer-based self-supervised learning (SSL) models also have fundamental differences: 1) they are trained on different modalities, some taking raw waveform as input while others using text; 2) they also vary in their training objectives, some using masked prediction while others using auto-regressive objects. Take Wav2Vec 2.0 and GPT-2 as examples, the former is a speech model trained on raw speech waveform using contrastive learning, while the latter is a language model trained on text corpus using next-word-prediction. Previous studies mainly focus on one specific type of models, either speech or text, and just compare the overall encoding prediction accuracy across models to find a better model. Therefore, it remains unclear what factors in the SSL features are contributing to the feature alignment between SSL models and brain. In other words, it is unknown if these different types of DNN models are correlated to the same aspects of neural activity underlying speech language processing. It is possible that the representation features from these DNN models of different modalities capture different components of the speech language, hence explaining different spatiotemporal factors in the population neural activity. On the other hand, it is also possible that different speech and language models converge to similar representations of speech language, and these converging factors also correlate to the neural activity underlying speech language processing.

This question has two folds: 1) if one particular SSL model could account for different aspects of neural dynamics during speech and language processing in the brain; 2) if SSL models of different modalities account for similar aspects in the neural activity during speech and language processing.

To delineate these alternatives and identify the critical factors driving the shared feature representation, we combined high-density electrocorticography (ECoG) recordings of the cortical network with different self-supervised deep neural network models of speech and language. We evaluated the representation similarity between self-supervised speech models, self-supervised language models, and the neural responses to naturalistic speech. Specifically, we built encoding models at single-word level^19^, instead of sentence-level which was more commonly used before^20–22^. This revealed more fine-grained temporal dynamics at the single-word level (i.e. a timecourse of prediction accuracy at different time offsets with respect to the word onsets). We also developed a CCA-based feature decomposition and encoding analysis^23^ to identify different levels of acoustic-linguistic information coding that drives these spatiotemporal dynamics.

## Results

### Overview

To investigate the neural activity underlying speech and language processing, we recorded ECoG signals from 15 native English speakers while they passively listened to 599 naturalistic English sentences^24^. Across all subjects, we identified 784 electrodes with significant speech responses in bilateral brain regions, including the superior temporal gyrus (STG), middle temporal gyrus (MTG), Heschl’s gyrus (HG), sensorimotor cortex (SMC), supramarginal gyrus (SMG), inferior frontal gyrus (IFG), middle frontal gyrus (MGF), etc. (see Methods and Supplement Fig. 1 for details).

We built neural encoding models to predict the spatiotemporal dynamics of neural activity using feature embeddings from pre-trained self-supervised learning models of different modalities (speech or text). For self-supervised learning models, we selected the representative speech model Wav2Vec2.0^10^ and the language model GPT-2^25^. These models took the raw audio waveforms and the corresponding transcribed text of the speech as input and transformed them into feature embeddings.

As shown in Figure 1, we analyzed ECoG neural activity within a 2-second window before and after the onset of each word. The analysis was conducted using a sliding window of 200 milliseconds with a step size of 20 milliseconds, calculating the average neural activity within each window. For each sliding window, we constructed linear deep neural encoding models. We used principal components analysis (PCA) to reduce the dimensionality of the word representations from the self-supervised learning models to 50 dimensions and employed these vectors to predict the average neural activity within each sliding window.

**Figure 1.**
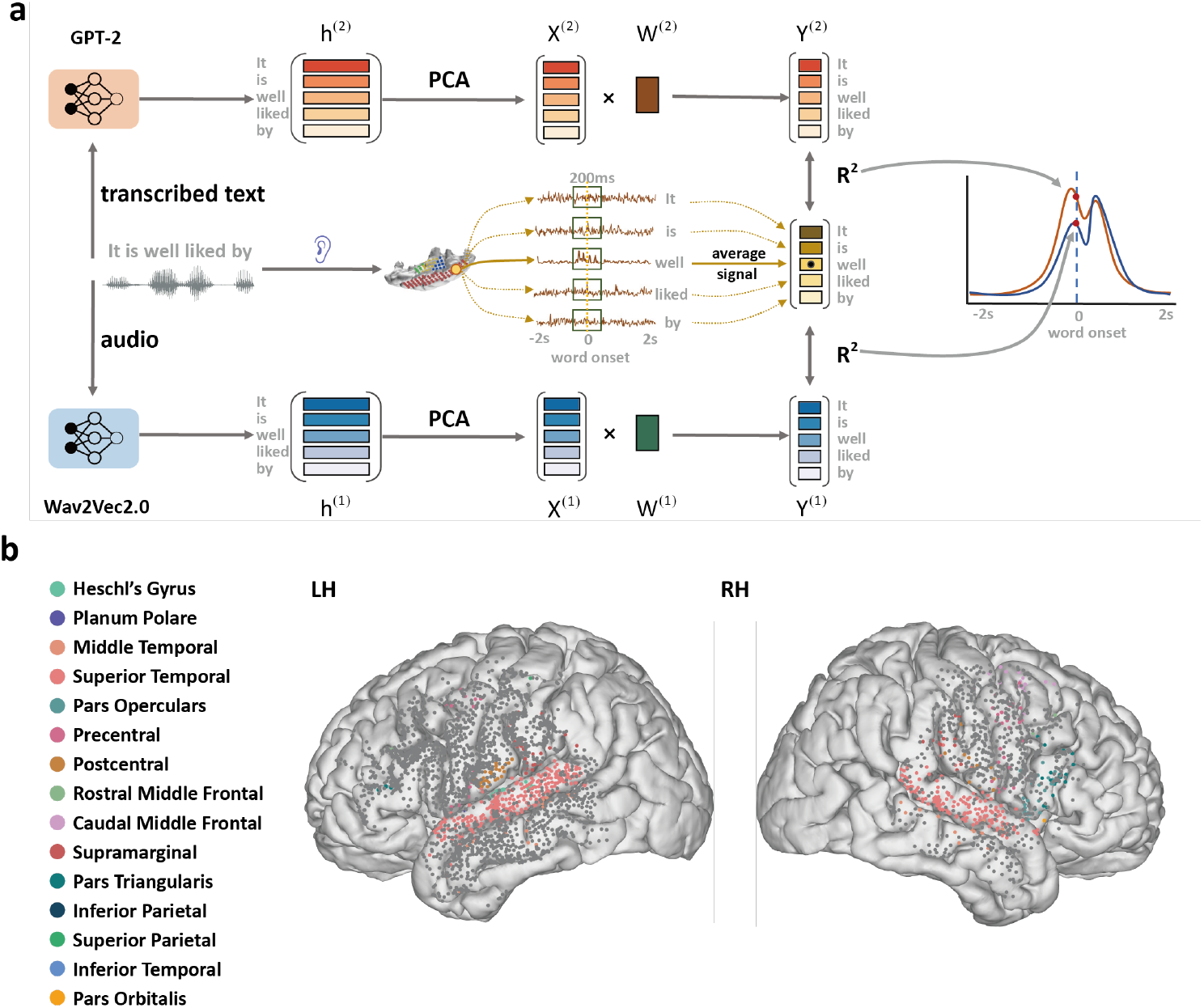
Neural encoding diagram and electrocorticography electrode distribution. **a)Diagram of word-level neural encoding using SSL models.** The neural activity of 15 epilepsy patients was recorded by Electrocorticography (ECoG) while listening to the TIMIT speech dataset. We captured the neural activity data 2 seconds before and after the onsets of each word. A sliding window of 200 milliseconds was set, moving in steps of 20 milliseconds. In each sliding window (lag), we established a linear encoding model to predict the neural activity. The neural activity within the lag was averaged to calculate the mean neural activity of all words. The audio and corresponding transcript from the TIMIT dataset were fed into Wav2Vec2.0 and GPT-2, respectively, to obtain word embedding vectors for speech and text. These vectors were then dimensionally reduced to 50 using Principal components analysis (PCA) and acted as the input to the linear encoding model to predict the neural activity. For each lag, we calculated the average *R*^2^ as encoding scores over speech responsive electrodes, allowing us to plot the timecourse of the encoding performance of Wav2Vec2.0 and GPT-2 before and after word onset. **b) Mapping of ECoG electrodes on a common brain surface**. Panel b illustrates the mapping of all ECoG electrodes onto a standard brain surface. Electrodes with significant responses to speech (colored electrodes) were selected for further analysis. Different colors correspond to distinct brain regions, while gray electrodes indicate a lack of significant response. The method for identifying significantly responsive electrodes is detailed in Method.

In addition to the data-driven SSL-based neural encoding models, we also built encoding models using three key components of speech representation for comparison^26^: static semantic information^27^, acoustic-phonetic information^28^, and contextual information^29^. Using features corresponding to these components, we predicted neural activity and subsequently compared these predictions with those from the self-supervised learning models. The prediction scores were obtained by calculating the squared Pearson correlation coefficient between the predicted and actual neural activity values. Each sliding window yielded a respective prediction score, allowing us to plot the temporal evolution of prediction scores within the 2-second window surrounding the word onsets.

### Both SSL models showed high neural prediction accuracy

First, we evaluated whether speech and text SSL models were able to predict neural activity with high accuracy. As illustrated in Figure 2a, throughout the 2-second interval surrounding word onset, the prediction scores of the speech self-supervised learning model Wav2Vec2.0 and the text self-supervised learning model GPT-2 were significantly higher than those of the three types of acoustic-linguistic features. This indicates that the variance in neural activity captured by the information encoded in the self-supervised learning models is substantially greater than that explained by individual linguistic features alone, highlighting the superiority of self-supervised learning models in modeling neural activity during speech perception.

**Figure 2.**
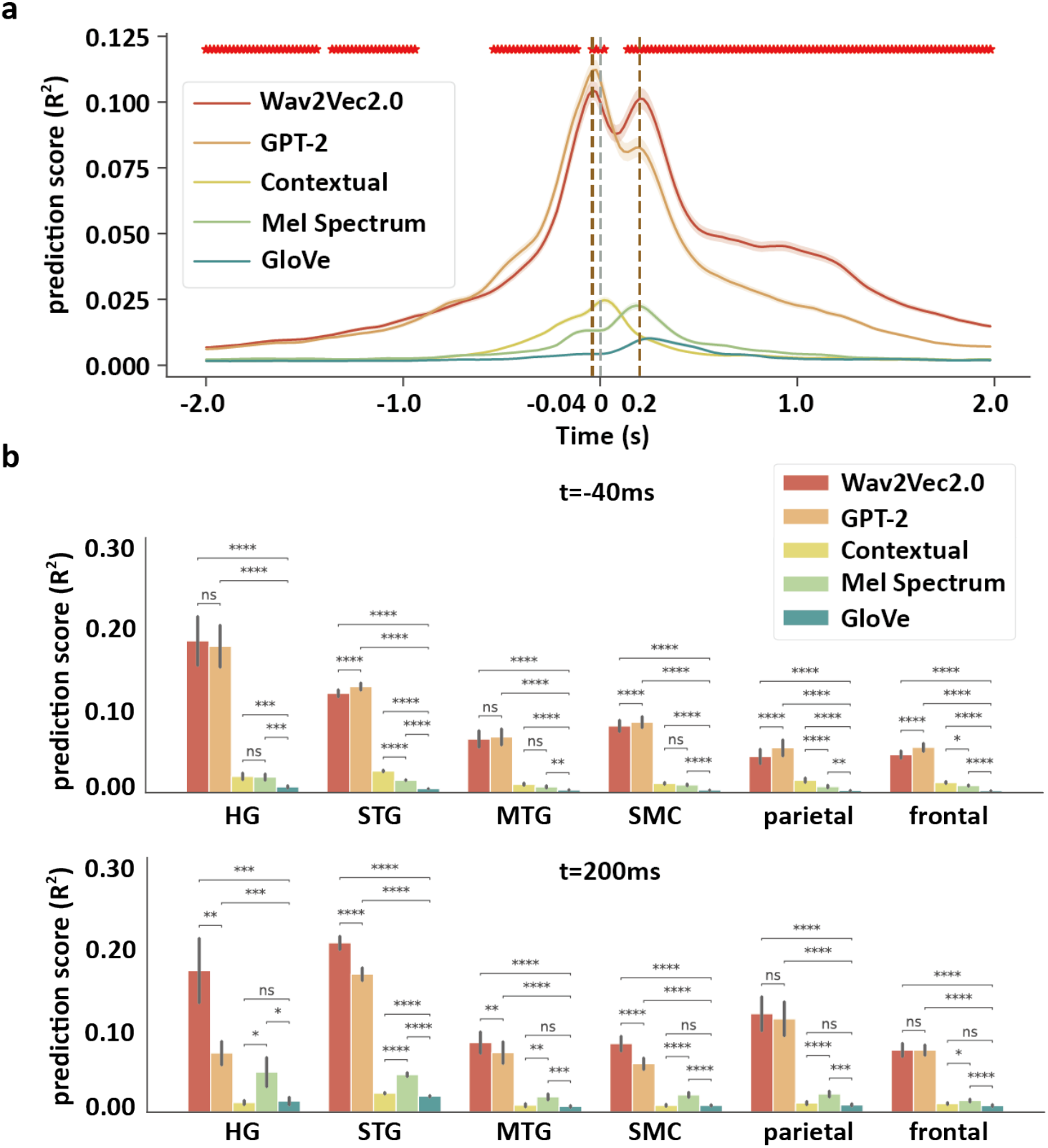
Temporal dynamics and regional distribution of neural encoding scores in speech processing. **a)** Brain encoding score of different features before and after word onset. ‘Mel Spectrum’ denotes the mel spectrum features, indicating acoustic-phonetic speech information. ‘Contextual’ denotes the contextual information^26^. ‘GloVe’ denotes static semantic information. Red asterisks indicate significant differences (Wilcoxon rank-sum, FDR correction, p < 0.05) in encoding scores between Wav2Vec2.0 and GPT-2 at the current time point. **b)** The average encoding scores of the SSL model in different brain regions 40ms before word onset (t=-40ms) and 200ms after word onset (t=200ms). **ns**: *p* > 0.05, *: *p* < 0.05, **: *p* < 0.01, ***: *p* < 0.001, and ****: *p* < 0.0001.

Moreover, the time course of the prediction scores for Wav2Vec2.0 and GPT-2 were remarkably similar, both peaking at 40 milliseconds before word onset (t=-40ms) and 200 milliseconds after word onset (t=200ms) (as shown by the brown dashed lines in Figure 2a). At 40 milliseconds before word onset (t=-40ms), the *R*^2^ for GPT-2 was slightly higher than that for Wav2Vec2.0 with significant difference d, with values of 0.112 and 0.104, respectively (Wilcoxon rank-sum, FDR correction, two-sided p = 0.041). Conversely, at 200 milliseconds after word onset, the trend reversed, with Wav2Vec2.0 achieving a higher *R*^2^ of 0.10 compared to GPT-2’s 0.08, with significant difference (Wilcoxon rank-sum, FDR correction, two-sided p = 6.949×10^-5^) .

The three baseline models also showed distinct temporal profile. The contextual feature model peaked at t=20ms around the word onset time with *R*^2^ = 0.025, while the acoustic model and the static semantic model peaked at t=180ms and t=260ms after the word onset, with *R*^2^ = 0.023 and *R*^2^ = 0.010 respectively.

### SSL models revealed spatiotemporal patterns of the entire language network

A finer-grain investigation into the spatial distribution of Wav2Vec2.0 and GPT-2 prediction accuracy revealed differential spatial patterns across the cortex. Specifically, we analyzed the prediction score in different regions-of-interests (ROI) at the two peak time points of the self-supervised learning models (Figure 2b). At t=-40 ms, GPT-2 showed significantly higher brain prediction accuracy than Wav2Vec2.0 in STG (t(503)= -10.367, p = 3.55e-22, two-sided), SMC (t(100)=-4.481, p=2.974e-05, FDR correction, two-sided), parietal (t(49)=-5.915, p=6.319e-07, FDR correction, two-sided) and frontal (t(75)=-7.461, p=3.767e-10, FDR correction, two-sided) cortices, and no significant difference in HG(t(13)=0.7796433995517917, p=0.462, FDR correction, two-sided) and MTG(t(38)=-1.730, p=0.100, FDR correction, two-sided) ; opposite pattern was found at t = 200ms, with Wav2Vec2.0 higher than GPT-2 in HG (t(13)=3.344, p=0.007, FDR correction, two-sided), STG(t(503)=11.767, p=1.882e-27, two-sided), MTG (t(38)=2.973, p=0.007, FDR correction, two-sided) and SMC(t(100)=6.717, p=4.121140764897982e-09, two-sided), and no difference in parietal(t(49)=0.872, p=0.436, FDR correction, two-sided) and frontal (t(75)=0.043, p=0.966, FDR correction, two-sided) cortices. These results suggest that GPT-2 better explains neural activities before the onset of each word at higher-order ROIs, such as STG and frontal areas, while Wav2Vec2.0 better accounts for neural activities after the onset of each word in areas associated with auditory perception, such as HG, SMC and STG.

Prediction scores of the three basic types of acoustic, phonetic and contextual-linguistic features also revealed consistent patterns (Figure 2b): at t=-40ms, higher prediction score of contextual features was found in STG, parietal and frontal cortices; at t=200ms, higher prediction score of acoustic (mel-spectrum) features was found in HG, STG, MTG, SMC, parietal and frontal areas. It is also worth pointing out that both models consistently showed higher prediction score than those of the three basic types of acoustic, phonetic and linguistic features (Figure 2b).

In summary, the distinct information encoded by SSL models may explain neural activity at different times and ROIs, potentially reflecting the spatiotemporal sequence of different types of information processing during speech perception across the large-scale language network.

### SSL models revealed fine-grained spatiotemporal dynamics in STG

In addition to the larger scale spatiotemporal dynamics at the ROI level, we investigate if SSL models reveal more detailed structures in particular ROIs. Along the anterior-posterior axis of STG, we found that the locations of high-scoring electrodes for Wav2Vec2.0 and GPT-2 change in time. At 40 milliseconds before word onset, the high-scoring electrodes of both models were primarily located in the middle part of STG, while at 200 milliseconds after word onset, the high-scoring electrodes were mainly in the posterior STG (Fig 2d).

To quantitatively compare the prediction scores of different neural populations in the STG, we divided the STG into 10 equidistant intervals along the y-axis (sagittal axis) and calculated the average prediction score for electrodes within each interval (Figure 3e). For Wav2Vec2.0, the prediction scores in middle STG (the interval y ∈ [0, 10]) at t=-40 ms than the prediction scores at t=200 ms. Conversely, in posterior STG (the interval y ∈ [−30, −10]) the prediction scores were higher at 200 milliseconds after word onset. Similar patterns were found for GPT-2, where the prediction scores in the interval from y ∈ [0, 15] (middle STG) were higher at t=-40 ms than t=200 ms, while in the interval around y = −20 (posterior STG), the opposite was observed.

**Figure 3:**
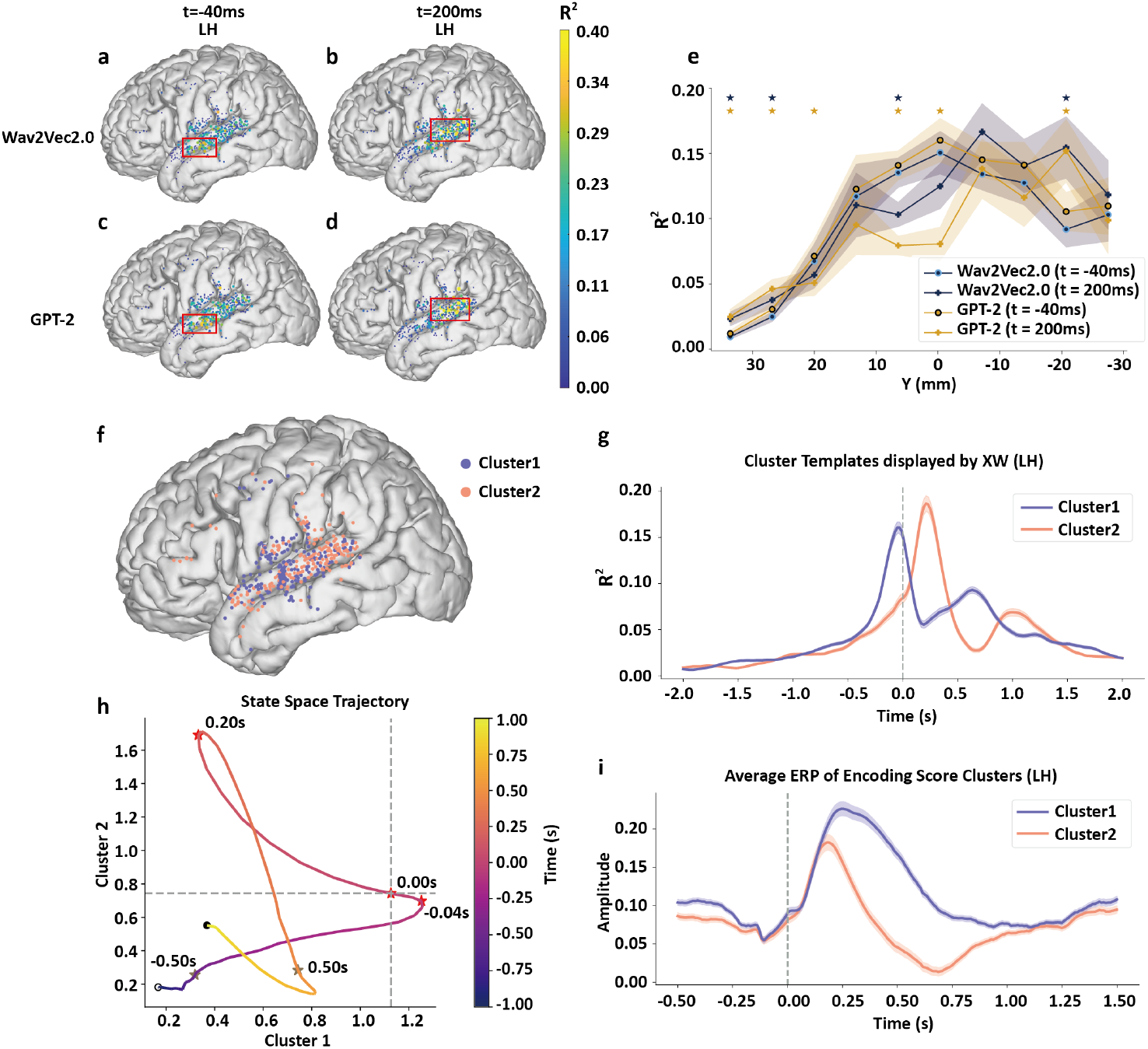
The transformer-based SSL models shows the ability to capture spatiotemporal dynamics in the brain. a to d) For the time before (t=-40ms) and after word onset (t=200ms), the neural populations with high encoding scores located in middle and posterior part in STG respectively. **e)** The distribution of encoding scores before and after word onset across electrodes on left hemisphere of all subjects. The line chart next to the brain surface map shows the average score over the electrodes in each bin on STG along the y-axis (divided into 10 bins). Blue stars above the graph indicate significant differences in Wav2Vec2.0’s performance between the two time points, while yellow stars indicate significant differences for GPT-2. *: *p* < 0.05. **f)** locations of electrodes from the two clusters plotted on a common brain surface. **g)** The averaged time courses of prediction score corresponding to the two clusters on left hemisphere. **h)** The state space trajectories of the entire neural response [-2s, 2s] around word onset, projected onto the two clusters. x-axis corresponds to the weights from Cluster 1 and y-axis corresponds to the weights from Cluster 2. **i)** the averaged speech-related neural response across all electrodes in each of the clusters, time 0 corresponds to speech onset.

Previous studies suggest that different subpopulations within STG account for complex lower- and higher-order information coding during speech and language perception^30^ . Furthermore, these subpopulations can be clustered based on their response profiles^31^ . We hypothesized that the pre- and post-word onset spatiotemporal peaks in Fig. 2 and 3a can be connected to and accounted by the dichotomy of functionally distinct populations. We employed convex non-negative matrix factorization to cluster the time series of prediction scores from SSL in the left hemisphere (see Methods). This clustering divided the electrodes into 2 categories(Fig. 3f). The two clusters coincided with the two peaks in the time course of prediction scores (Fig. 3g): Cluster 1 (marked in orange) had a primary peak of prediction score at around 40ms before word onset and a later secondary peak 500ms after word onset; Cluster 2 (marked in purple) had primary peak at 200 ms after word onset. Moreover, the two clusters showed distinct onset speech response profiles: Cluster 1 mainly demonstrated a more sustained speech onset response, extending to 500 ms after onset (Fig. 3i); Cluster 2 had a more transient onset response peaked around 200 ms after speech onset (Fig. 3i). Anatomically, Cluster 1 mainly located in the middle and anterior STG, ventral SMC, as well as posterior middle frontal gyrus (MFG); Cluster 2 mainly located in the posterior STG and the temporal parietal junction.

From a state space dynamic point of view (Fig. 3h), the population activity was firstly driven by Cluster 1 before word onset and peaked at 40ms before onset, then the dynamics rapidly switched to Cluster 2 and peaked around 200ms after word onset, and gradually shifted back to Cluster 1 at around 500ms after onset. This state space circular trajectory suggests that SSL models are able to capture single-word level dynamics during continuous speech perception.

In summary, clustering prediction scores from self-supervised learning models reveals distinct activity patterns of different neural populations in the speech language cortical network. These results indicate the relationship between the prediction scores of self-supervised learning models and the complex spatiotemporal patterns in the cortex. This demonstrates the ability of Transformer-based self-supervised learning models to capture diverse spatiotemporal patterns in the brain corresponding to speech and language processing. In other words, the same SSL model could learn diverse representation that accounts for different aspects of the speech language network. Furthermore, word-level dynamics can be revealed using SSL-model-based encoding models.

### Shared acoustic-phonetic and contextual information explained the temporal dynamics in brain

Previous results suggest that the same SSL model can elucidate the processing sequence of different types of information during speech perception. Next, we ask whether SSL models of different modalities explain the same aspects in the spatiotemporal neural dynamics. Qualitatively examining the predicted neural activity by Wav2Vec2.0 and GPT-2 suggested that (Supplement Fig. 3) the predictions from both SSL models were able to capture similar patterns of neural activity. We calculated the squared Pearson correlation coefficient between the predicted neural activities of the two models as a metric of their similarity (Figure 4d, denoted as W2V2_GPT2). Additionally, we computed similarity scores between the neural activity in left hemisphere predicted by Wav2Vec2.0, GPT-2, and Mel spectrograms as baselines for comparison (Figure 4d, denoted as GPT2_Mel and W2V2_Mel). The neural activity predictions from Wav2Vec2.0 and GPT-2 were highly similar, with similarity scores (*R*^2^) exceeding 0.35 across various brain regions associated with the language network, significantly higher than those of the baseline models. Specifically, the similarity scores in STG between Wav2Vec2.0 and GPT-2 predictions were significantly higher than those between Wav2Vec2.0 and Mel spectrogram predictions (t(699)= 45.948, p= 2.029e-212, FDR correction, two-sided) and those between GPT-2 and Mel spectrogram predictions (t(699)=54.936, p =1.386e-254, FDR correction, two-sided).

**Figure 4:**
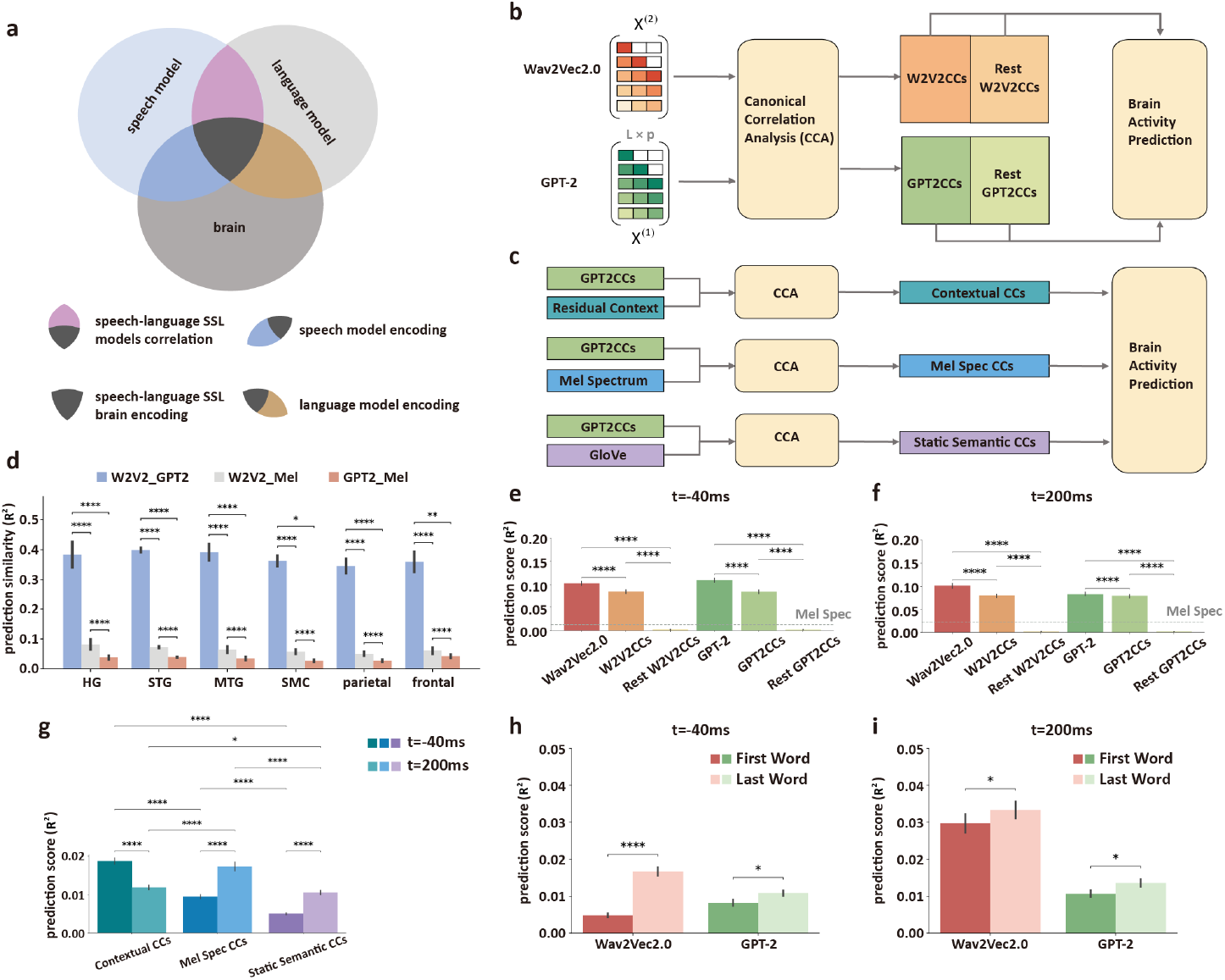
Decomposition of the SSL-based feature encoding analysis using Canonical Correlation Analysis (CCA). **a)** Diagrams of the variance decomposition among speech model, language model and the brain. **b)** Extracting shared representations between GPT-2 and Wav2Vec2.0 using CCA and predicting the neural response using the extracted CC components (GPT2CCs, W2v2CCs); **c)** Decomposing the shared CC components into contextual, static semantic and acoustic-phonetic information and predicting the neural response using the decomposed CCs. d) Prediction similarity (R^2^) between models across brain regions: HG (Heschl’s Gyrus), STG (Superior Temporal Gyrus), MTG (Middle Temporal Gyrus), SMC (Sensorimotor Cortex), parietal, and frontal. W2V2_GPT2 indicates similarity between Wav2Vec2.0 and GPT-2, W2V2_Mel between Wav2Vec2.0 and Mel Spectrum, and GPT2_Mel between GPT-2 and Mel Spectrum. Similarity was calculated for predictions at t=-40ms and t=200ms, averaged across these time points. The result demonstrates the high similarity of neural activities and the shared components between Wav2Vec2.0 and GPT-2. **e and f)** Encoding score of Wav2Vec2.0, GPT2 and their shared components before (t=-40ms) and after (t=200ms) word onset. **g)** The encoding score of the decomposed components: contextual (Contextual CCs), static semantic (Static sematic CCs), and acoustic-phonetic (Mel Spec CCs) information in SSL features before (t=-40ms) and after (t=200ms) word onset. The p-values of the significance test were obtained through paired t-test and FDR correction. **h) and i)** Encoding score of Wav2Vec2.0 and GPT-2 for the first and last words of sentences at before (t=-40ms) and after (t=200ms) word onset. **ns**: *p* > 0.05, *: *p* < 0.05, **: *p* < 0.01, ***: *p* < 0.001, and ****: *p* < 0.0001.

The above results indicate a high degree of similarity in word-level neural activity predictions by the two types of SSL models. To further investigate whether this similarity arises from shared or unique information of these two models, we employed canonical correlation analysis (CCA) to decompose the model representation into shared and unique information between the two feature spaces (see Methods and Figure 4a). In particular, CCA identifies and measures the relationships between the two feature spaces by finding linear combinations of the variables in each model’s feature space that are maximally correlated with each other, resulting in a sorted list of canonical components (CCs). We extracted the shared information between the word feature matrices of Wav2Vec2.0 and GPT-2 using top 50 CCs (Figure 4b, denoted as W2v2CCs and GPT2CCs), and the unique information of each SSL model, using the remaining CCs (Figure 4b, denoted as Rest W2v2CCs and Rest GPT2CCs). These features were then used to build different neural encoding models and to predict neural activity. We compared the prediction scores of the full SSL model features, the shared information features, and unique information features at the two peak times: 40 milliseconds before (t=-40ms) and 200 milliseconds after (t=200ms) word onset. For both time points, the prediction scores from the shared information were close to those of the original full features (Wav2Vec2.0 and GPT-2)(Fig. 4e and 4f, we denoted prediction score as “ps”, ps(t=-40ms, Wav2Vec2.0)=0.103 vs ps(t=-40ms, W2V2CCs)=0.085, t(490)=23.291, p=3.704e-81, ps(t=200ms, Wav2Vec2.0)=0.102 vs ps(t=200ms, W2V2CCs)=0.079, t(490) = 20.209, p=5.049e-66; ps(t=-40ms, GPT-2)=0.110 vs ps(t=-40ms, GPT2CCs)=0.085, t(490)= 31.867, p=9.676e-121; ps(t=200ms, GPT-2)=0.084 vs ps(t=-40ms, GPT2CCs)=0.078, t(490) = 5.834, p=9.862e-09; paired t-test, FDR correction, two sided) and significantly higher than those from unique information (Fig. 4e and 4f, ps(t=-40ms, Wav2Vec2.0)=0.103 vs ps(t=-40ms, Rest W2V2CCs)=0.002, t(490) = 23.730, p=3.832e-83, ps(t=200ms, Wav2Vec2.0)=0.102 vs ps(t=200ms, Rest W2V2CCs)=0.002, t(490) = 20.607, p=1.236e-67; ps(t=-40ms, GPT-2)=0.110 vs ps(t=-40ms, Rest GPT2CCs)=0.002, t(490) = 24.823, p=3.330e-88; ps(t=200ms, GPT-2)=0.084 vs ps(t=200ms, Rest GPT2CCs)=0.0019, t(490) = 17.797, p=1.014e-54; paired t-test, FDR correction, two sided) . This indicates that the shared information between Wav2Vec2.0 and GPT-2 accounted for the majority of the variance in neural activity.

Next, we aimed to understand the specific linguistic information encoded in the shared information. We analyzed three aspects of speech representation^26^: form (acoustic-phonetic features), meaning (semantics), and structural/contextual information. To capture these, we employed the mel spectrum^28^ for form, GloVe^27^ embeddings for meaning, and residual context embeddings^29^ for structural context . Similar to the previous CCA-based approach, we applied a second-level of CCA. In particular, we used CCA to extract canonical variables between the shared information (the word feature matrix GPT2CCs) and the three key components of speech representation, incorporating these into the neural encoding models (Figure 4g). Figure 4g show the prediction scores for context, static semantics, and acoustic-phonetic information within the shared information at 40 milliseconds before (t=-40ms) and 200 milliseconds after (t=200ms) word onset. At 40 milliseconds before word onset, the prediction score for contextual information (*R*^2^ =0.019) was significantly higher than those for the other two features (we denoted prediction score as “ps”, ps(t=-40ms, Contextual CCs)=0.019 vs ps(t=-40ms, Mel Spec CCs)=0.009, t(490)=13.952, p=1.717e-37; ps(t=-40ms, Contextual CCs)=0.019 vs ps(t=-40ms, Static Semantic CCs)=0.005, t(490) = 20.005, p=1.598e-65; paired t-test, FDR correction, two sided). This indicates that contextual information explained most of the variance in neural activity before word onset. At 200 milliseconds after word onset, the prediction scores changed, with the Mel spectrogram (*R*^2^=0.0017) surpassing the other two features (we denoted prediction score as “ps”, ps(t=200ms, Mel Spec CCs)=0.017 vs ps(t=200ms, Contextual CCs)=0.012, t(490)=-6.257, p=8.578e-10; ps(t=200ms, Mel Spec CCs)=0.017 vs ps(t=200ms, Static Semantic CCs)=0.011, t(490)=-7.876, p=2.1952e-14; paired t-test, FDR correction, two sided). This suggests that acoustic-phonetic information explained most of the variance in neural activity after word onset.

To further dissociate and validate the neural coding of these distinct sets of information, we evaluated the performance of encoding models at different positions of each sentence, where contextual information varied. Since we presented the sentences in random order, the amount of contextual information distributed unevenly at different temporal landmarks: the last word should contain more contextual information compared to the initial word of each sentence, particularly for the t=-40ms timestamp. We found that for t=-40 ms before word onsets, both models showed higher encoding performance for the last word of each sentence, compared to the first word (ps(t=-40ms, Wav2Vec2.0, First Word)=0.005 vs ps(t=-40ms, Wav2Vec2.0, Last Word)=0.017, t(490)=-9.164, p=1.377e-18; ps(t=-40ms, GPT-2, First Word)=0.008 vs ps(t=-40ms, GPT-2, Last Word)=0.011, t(490)=-2.369, p=0.018; paired t-test, FDR correction, two sided). On the other hand, the bottom-up processing driven by acoustic-phonetic information should have little dependency on the temporal position in each sentence. At t=-200ms, although the encoding score for the last word remains significantly higher than that of the first word, the gap between them has narrowed compared to the results at t=-40ms. This suggests that the contextual advantage for last words in sentences is less pronounced and acoustic-phonetic information is the main driver of neural responses at this time point (ps(t=200ms, Wav2Vec2.0, First Word)=0.030 vs ps(t=200ms, Wav2Vec2.0, Last Word)=0.033, t(490)=-2.454, p=0.015; ps(t=200ms, GPT-2, First Word)=0.011 vs ps(t=200ms, GPT-2, Last Word)=0.014, t(490)=-2.430, p=0.015; paired t-test, FDR correction, two sided) .

In summary, the results indicate that the variance in neural activity explained by different types of information encoded in the features of SSL models changes over time. Contextual information mainly explains the variance in neural activity before word onset, while acoustic-phonetic information explains most of the variance after word onset. This may reflect a temporal sequence in the brain’s processing of linguistic information: before hearing a word, the brain primarily processes contextual information and predict the incoming speech^19^ ; after hearing the word, it is mainly driven by bottom-up acoustic-phonetic information.

## Discussion

In this study, we explored the spatiotemporal dynamic relationship between self-supervised learning models and neural activity during speech perception. Using ECoG signals from 15 subjects during speech perception, we demonstrated the correspondence between self-supervised learning speech and text models and the brain in terms of spatial, temporal, and neural activity patterns, as well as information encoding. This indicates the powerful modeling capability of self-supervised learning models for neural activity. Specifically, our contributions include three main points.

This research theme holds significant importance in the current fields of neuroscience and natural language processing. With the continuous development of self-supervised learning models, researchers have begun to use self-supervised learning models to study the language processing mechanisms of the human brain^16,18,19,32–34^. These studies have shown that self-supervised learning models can capture neural activity patterns in the human brain during language processing. However, our understanding of the spatiotemporal dynamic relationship between self-supervised learning models and neural activity remains limited. Previous research has often demonstrated correlations between features of a particular neural network layer and neural activity in specific parts of the brain’s language network from a macro perspective, but there is a lack of detailed descriptions and characterizations of the different spatiotemporal components in neural activities of the language network.

First, we demonstrated that self-supervised learning models can comprehensively describe the dynamic neural encoding process of speech and language perception. Beyond validating that the feature embeddings from the speech model and the text model both predict neural activity well, we constructed word-level neural encoding models at different time-offsets relative to the word onsets and revealed distinct temporal profiles of prediction accuracy with the two models. The results indicated that both self-supervised learning models achieved high prediction performance at 40 milliseconds before and 200 milliseconds after word onset. These two temporal landmarks corresponded to the representation of dynamic contextual information and the acoustic-phonetic information respectively, distributed across the broad speech language processing network in the temporal auditory areas (HG and STG), parietal lobe (supramarginal), and the frontal areas (SMC, IFG, and MFG). Previous studies in the literature have shown that embeddings in the SSL speech model mapped to different neural populations spatially^32^, and embeddings in the SSL language model correlated to the temporal dynamics in the neural population ^18^. Here we provide a unified dynamic view that the same SSL model can reveal the spatiotemporal across the brain^35,36^. This is one step forward towards more general computational models that account for different cognitive aspects of speech and language processing^30,37^ . Together with several recent studies^16,18,32,38^, we show that self-supervised models are able to characterize the broad spatiotemporal dynamics across the speech and language processing network in the brain.

Our results also revealed fine-grained population dynamics in high-order speech cortex, such as STG. Previous studies have investigated the high-order speech cortex from the perspectives of both local neural coding and large-scale population dynamics. On one hand, distinct local populations in STG encode rich and complex acoustic-phonetic information during speech perception^30,37,39^ . On the other hand, population dynamics in the STG track the temporal landmarks and context in speech, such as the onsets of a sentence^31^, or word-level cues^40,41^ . We show that under a unified framework based on SSL models, both the local coding properties, and the populational word-level and sentential level context representation dynamics can be revealed and analyzed.

Finally, we discovered that self-supervised learning models learned shared information from different modalities—speech and text—and that different linguistic information within this shared information contributed to the variance in neural activity before and after word onset. Using CCA, we analyzed the variance in word-level neural activity attributed to different linguistic components encoded in the shared information of speech and text self-supervised learning models. The study found that 40 milliseconds before word onset, contextual information contributed the most variance, whereas 200 milliseconds after word onset, acoustic-phonetic information contributed the most variance. This suggests that during speech perception, the brain utilizes linguistic cues from different information sources, with different linguistic information playing dominant roles in different sliding windows. This finding is consistent with existing behavioral and electrophysiological research, which indicates that contextual information can influence early speech processing^42^, while acoustic-phonetic information is processed at later stages^43^. Additionally, the study found that the bottom-up feedforward process reflected by acoustic-phonetic information interacts with the neural population’s internal recurrent process reflected by contextual information in the STG, collectively influencing the speech perception process. This reveals that the same neural circuits may be involved in integrating multiple levels of information during speech perception, similar to findings in visual perception^44^, suggesting that there may be similar computational mechanisms in different modalities of brain perception.

In summary, our work used self-supervised learning models of both speech and text modalities to construct deep neural encoding models, uncovering the correspondence between the linguistic information encoded in self-supervised learning models and the spatiotemporal dynamics of neural activity during speech perception. By leveraging the spatiotemporal correspondence between self-supervised learning models and neural activity, this research further supports existing findings on the computational principles of brain language processing, such as the correspondence between functional characteristics and brain anatomical structures^45^, and the interaction of bottom-up and top-down processes during speech perception^37^. The innovative aspect of this study lies in performing clustering analysis on the prediction scores of neural activity by self-supervised learning models, dividing neural populations based on temporal dynamic characteristics from a functional perspective, and discovering that this functional division corresponds to two distinct neural populations in spatial anatomical structures. This finding indicates that self-supervised learning models have learned the spatiotemporal dynamic patterns of neural activity in the human brain during speech perception, reflecting the fine-grained characterization of the neural computation process of speech perception by self-supervised learning models. This provides important reference for constructing more accurate and comprehensive neural computational models of speech perception.

## Materials and Methods

### ECoG dataset

We conducted ECoG recordings on 15 native English speakers while they listened to 599 sentences from the TIMIT corpus^46^. ECoG offers high spatiotemporal resolution and signal-to-noise ratio compared to non-invasive methods like fMRI or EEG. Electrode grids were implanted during epilepsy treatment, covering the cortical language network in temporal, parietal, and frontal lobes (Supplement, Fig. 1. We analyzed 784 speech-responsive electrodes out of a total of 4388, primarily located in the superior temporal gyrus (STG), sensorimotor cortex (SMC), middle temporal gyrus (MTG) and supramarginal gyrus (SMG).

For each electrode, the local field potential underwent amplification and was recorded at a frequency of 3,052 Hz. Visual inspection was conducted on the raw voltage waveforms, leading to the removal of channels that exhibited signal variations too minor to distinguish from noise or consistent epileptiform activities. For the remaining channels, segments with electrical or movement-related distortions were identified manually and omitted. The signal underwent a notch filtering process to eliminate line noise, set at frequencies of 60, 120, and 180 Hz. This was followed by a re-referencing to the averaged common signal across channels connected to the same preamplifier. Using the Hilbert transform, the analytic amplitude for eight Gaussian filters, with a central frequency range of 70–150 Hz, was calculated. The average analytic amplitude across these eight frequency bands was then used to represent the high-gamma signal. This signal was subsequently downsampled to 100 Hz. The process of recording was structured into intervals, each lasting about 5 minutes, during which the high-gamma signal underwent z-score normalization across each recording segment.

### Speech responsive electrodes selection

We compared the actual neural activity 2 seconds before and after the onset of each sentence with the threshold derived from randomly sampled neural activity. If the neural activity of a particular electrode significantly exceeded the threshold, we considered that electrode to be significantly responsive to the speech stimulus.

Specifically, for the neural signals of each electrode, we conducted 1000 random samplings. Each sampling involved selecting 599 random sentence onsets (to match the 599 sentences in the TIMIT dataset) and extracting the neural data within a 4-second sliding window, spanning 2 seconds before and after each random time point (resulting in a total of 400 data points, corresponding to different time instances). For each randomly sampled time point, we computed the mean and standard deviation across all samplings. This allowed us to determine the mean and standard deviation for each time point within the 2-second window around the random time points. We set the threshold for the current time point as the mean plus five times the standard deviation.

For the onset of each sentence, we compared the actual neural activity with this threshold. If there were 10 consecutive data points (corresponding to 100 milliseconds of neural activity) that exceeded the threshold, we considered the electrode to be significantly responsive to the speech stimulus.

### Encoding Analysis

Figure 1 illustrates the overall workflow of the encoding analysis. First, we extracted the intermediate layer activations of self-supervised learning models corresponding to the TIMIT dataset. The audio waveforms and corresponding transcriptions from the TIMIT dataset were used as inputs to the speech self-supervised learning model Wav2Vec2.0 and the text self-supervised learning model GPT-2, respectively, generating activation sequences h = [h(1), …, h(L)]^*T*^ ∈ R^L×d^, where d represents the embedding dimension (768 for Wav2Vec2.0 and 1600 for GPT-2 XL), and L = 3543 represents the number of non-deduplicated words in the audio transcription, arranged sequentially according to their presentation order. To reduce computational complexity, we applied Principal Component Analysis (PCA) to reduce the word embeddings to 50 dimensions, denoted as d_p_. Thus, the input to the neural encoding model is 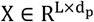.

For the neural activity within 2 seconds before and after each word onset (totaling 4 seconds), we calculated the average neural activity in sliding windows of 200 milliseconds (with a 20-millisecond step size). For each sliding window, we used the word feature matrix composed of word embeddings as the input to the neural encoding model to predict the average neural activity for all words in that window. For the t_1_-th sliding window, the corresponding neural activity matrix is denoted as 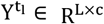, where 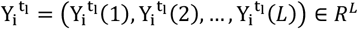, and 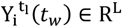 represents the average ECoG signal for the i-th electrode on the t_w_-th word. Here, i = 1, …, c, where c represents the total number of electrodes with significant responses to speech, and t_1_ denotes the current sliding window index, with t_1_ = 1, …, N_lags_, where N_lags_ = 200 in this study. For each self-supervised learning model, the prediction score for each electrode is the highest prediction score across all layers of the model.

For each sliding window t_1_ and each independent electrode i, we fit an L2-regularized linear regression model to predict neural activity:

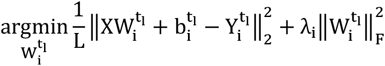

where i = 1, …, c, 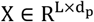 denotes the word feature matrix, 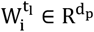, and 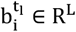 represent the weights and biases of the linear regression model for the i-th electrode in the t_1_-th sliding window, respectively. λ_i_ denotes the L2 regularization coefficient for the i-th electrode.

We obtained the prediction scores for each time point around word onset by calculating the R^2 between the predicted and actual neural responses for each sliding window. Specifically, for each ECoG electrode i, the prediction score of the self-supervised learning model is the optimal R^2^ across all Transformer layers. For each electrode i (i = 1, …, c) in the t_1_-th sliding window, the R^2^ is calculated as the squared Pearson correlation coefficient between the predicted neural activity 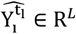 and the actual values 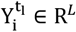:

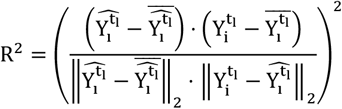

This approach provides a comprehensive analysis of how well the self-supervised learning models predict neural activity across different time points and spatial locations in the brain.

### Clustering Analysis of Prediction Scores Based on Convex Non-negative Matrix Factorization (CNMF)

To analyze the prediction scores and activity patterns of different neural populations, we employ convex non-negative matrix factorization (CNMF) to cluster the prediction scores from the Wav2Vec2.0 model for each ECoG electrode. This approach allows us to categorize different neural populations based on the patterns of their prediction scores. Specifically, let the prediction score matrix be 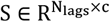, where the matrix element S_ij_ represents the prediction score of the j-th electrode in the i-th sliding window, N_lags_ represents the total number of sliding windows, and c represents the number of electrodes. We applied CNMF to decompose S:

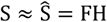

where 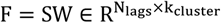, with each row representing the weighted prediction score sequence of each cluster, reflecting the temporal activity patterns captured by each category. The matrix 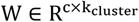 represents the weights of the prediction score sequences for each electrode. The matrix 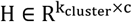 has each column representing the weight of an electrode in the k_cluster_ clusters, where the row index with the maximum value in each column indicates the cluster to which the electrode belongs.

To estimate the optimal number of clusters, we calculated the percentage of variance explained by projecting the prediction activities onto their corresponding clusters using the following formula:

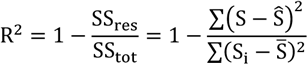

where SS_res_ is the residual sum of squares, capturing the squared differences between the CNMF predictions and the actual values, representing the unexplained variability in the data due to the decomposition. SS_tot_ represents the total sum of squares, measuring the total variance of the actual data from its mean, indicating the overall variability in the data. Thus, the R^2^ in the above formula represents the proportion of the total variance in the dependent variable explained by the independent variables in the CNMF. When k = 2, the R^2^ value for all matrices consistently remains above 0.9, so we set the number of clusters to 2.

For visualizing the state space trajectories, we projected all electrode data onto the W matrix obtained from the previous CNMF method. For instance, after applying CNMF to the prediction electrode matrix of the left hemisphere (with dimensions 200 × 491), we obtained W ∈ R^491×2^ and H ∈ R^2×491^. The H matrix provides classification factors for the 491 significant electrodes, with each column containing two rows of data. If the value in the upper row exceeds that in the lower row, the electrode is assigned to the first cluster; otherwise, it is assigned to the second cluster. Subsequently, based on the classification results of H, the data S and matrix W were weighted according to the two electrode clusters, resulting in S_1_W_1_ for the first cluster and S_2_W_2_ for the second cluster, where S_1_ and S_2_ represent the data of the two electrode clusters, and W_1_ and W_2_ represent only the first and second columns of W, respectively. S_1_W_1_ and S_2_W_2_ constitute the word trajectories in the state space.

### Shared Representation Extraction Based on Canonical Correlation Analysis

We aim to explore how the shared and unique information from Wav2Vec2.0 and GPT-2 each contributes to the explainable variance in neural activity. To achieve this, we used CCA to extract the canonical variables between the feature spaces of Wav2Vec2.0 and GPT-2, thereby obtaining their shared and unique information. CCA^47^ is a multivariate statistical method used to explore the correlations between two sets of variables. The fundamental idea of CCA is to find linear combinations of the two sets of variables such that the correlation between these linear combinations is maximized, revealing the underlying association patterns between the variables.

Let 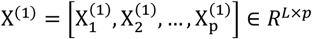 and 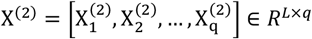 represent the feature matrices of Wav2Vec2.0 and GPT-2 on the TIMIT dataset, respectively. The goal of CCA is to find linear combinations of the column vectors of these feature matrices, 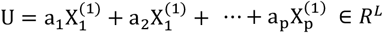 and 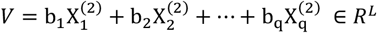, such that the correlation coefficient between U and V is maximized. U and V is a pair of the canonical variables, with a = (a_1_, a_2_, …, a_p_)^T^ ∈ *R*^*p*^ and b = (b_1_, b_2_, …, b_q_)^T^ ∈ *R*^*q*^ representing the weight vector for the two sets of variables, respectively.

The optimization objective can be expressed mathematically as:

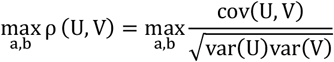

where ρ(U, V) denotes the correlation coefficient between U and V, cov(U, V) represents the covariance between U and V, and var(U) and var(V) represent the variances of U and V, respectively.

We consider the first 50 pairs of canonical variables (CCs) obtained through CCA to sufficiently capture the shared information between Wav2Vec2.0 and GPT-2, while the remaining CCs represent the unique information of each model. These canonical variables are then used as inputs to the neural encoding models to predict neural activity. By evaluating the prediction scores of shared and unique information on neural activity, we can compare their contributions to the explainable variance in neural activity.

### Disserting Specific Information in Shared Components

Linguistic theory posits three key dimensions within speech-language: acoustic-phonetic information, static semantics, and structural/contextual information^26^. We further modeled these three aspects as three models and used them to dissect the shared representations between SSLs. Inspired by Pasad et al^48^, we applied CCA to analyze the static semantic information and acoustic-phonetic information encoded in the shared representation between SSLs, probing with GloVe and mel spectrum respectively. For static semantics, we employed GloVe embeddings^27^, tailored to capture semantic meaning independently of context. To represent acoustic-phonetic features, we utilized the mel spectrum.

Contextual information was captured through residual context embeddings, as introduced by Toneva et al^29^. This approach extracted contextual information from GPT-2 by regressing out first-layer embedding features from the embeddings of the intermediate layer, resulting in contextual representations excluding lexical-level features from the first layer. Specifically, we denote h_k_(t − 1) ∈ *R*^*d*^ as the activation of the k’th transformer layer for the (t − 1)’th word in SSL model.

To obtain the contextual representations, we first used ridge regression to learn projection onto h_k_(t − 1). The model’s input is a concatenated matrix H_t_ = [h_0_(t − 24), h_0_(t − 23), …, h_0_(t)] ∈ R^d×25^, where each h_0_ is a word vector from layer 0 at the current and the preceding 24 words.

Let g(x) denote the ridge regression model that projects onto h_k_(t − 1). We use ct_k_ (t − 1)to represent the contextual information at time t − 1:

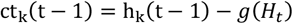

Where t = 1, .., L, and k set to 9 for GPT-2, which corresponding to the optimal layer decoder8 in GPT-2.

## Data availability

The raw datasets supporting the current study have not been deposited in a public repository because it contains personally identifiable patient information, but are available in an anonymized form from E.F.C. on reasonable request.

## Code availability

The completely developed code that operates on the full dataset will be made available from the authors upon reasonable request. Source code that implements the core neural encoding algorithm and the further analysis can be found at https://github.com/chenperry/brain_encoding.

## Author contributions

Conceptualization: P.C. and Y.L.; methodology: P.C., L.H., Y.L.; software: P.C., S.X., L.H., and Y.L.; formal analysis: P.C. and Y.L.; resources: E.F.C.; writing—original draft: P.C.; writing—review and editing: P.C., S.X., and Y.L.; supervision: Y.L.; project administration: Y.L.

## Acknowledgements

This work is supported by the National Natural Science Foundation of China (32371154, Y.L.), Shanghai Rising-Star Program (24QA2705500, Y.L.), Shanghai Pujiang Program (22PJ1410500, Y.L.), and the Lingang Laboratory (LG-GG-202402-06-08, Y.L.). The computations in this research are supported by the HPC Platform of ShanghaiTech University.

