## Supplementary Information for "Convergent representations and spatiotemporal dynamics of speech and language in brain and deep neural networks"

Chen *et al.*

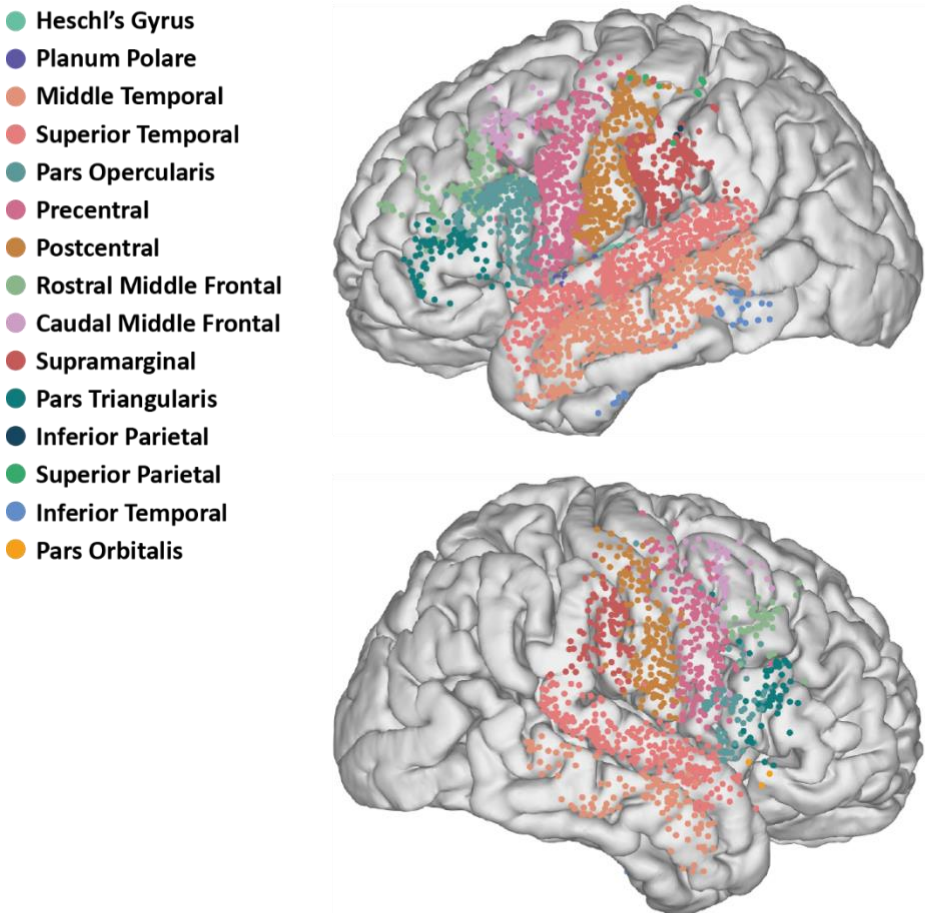


**Supplementary Figure 1. Spatial distribution of all ECoG electrodes.** The upper part of the image represents the left hemisphere, and the lower part represents the right hemisphere. There are a total of 4,388 electrodes covering regions associated with the language network. We identified 784 electrodes that showed significant responses to speech stimuli in bilateral brain regions.(see Methods and Figure 1 for details).

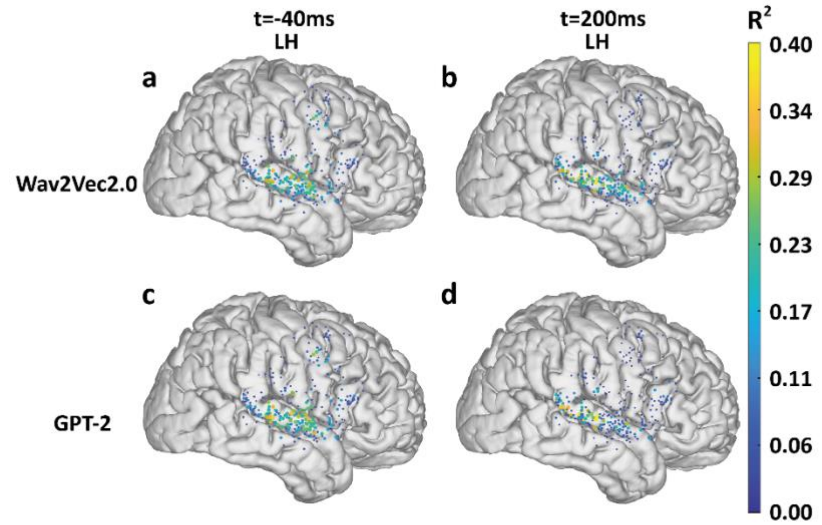


**Supplementary Figure 2. Encoding score distribution on right hemisphere.** For the time before (t=-40ms) and after word onset (t=200ms), the neural populations with high encoding scores located in middle and posterior part in STG respectively.

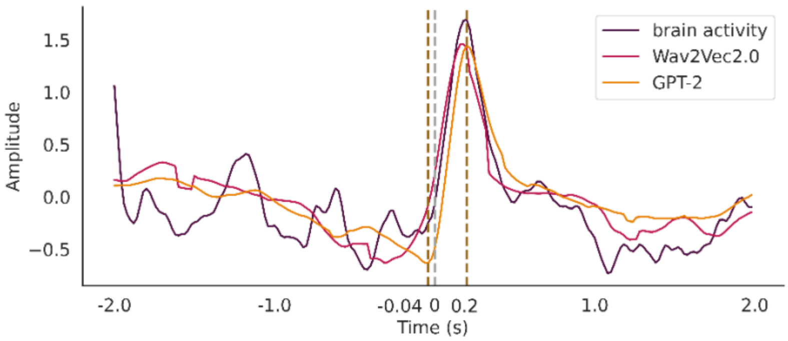


**Supplementary Figure 3. Actual and predicted brain activity of an example word.** The figure compares the actual brain activity with the predicted brain activity for an example word, using features extracted from speech model Wav2Vec2.0 and language model GPT-2. The close resemblance between the actual and predicted curves indicates that features from both models can effectively capture neural activity patterns during language processing, which highlights the potential of using advanced language models to understand and predict brain responses associated with speech and language comprehension.

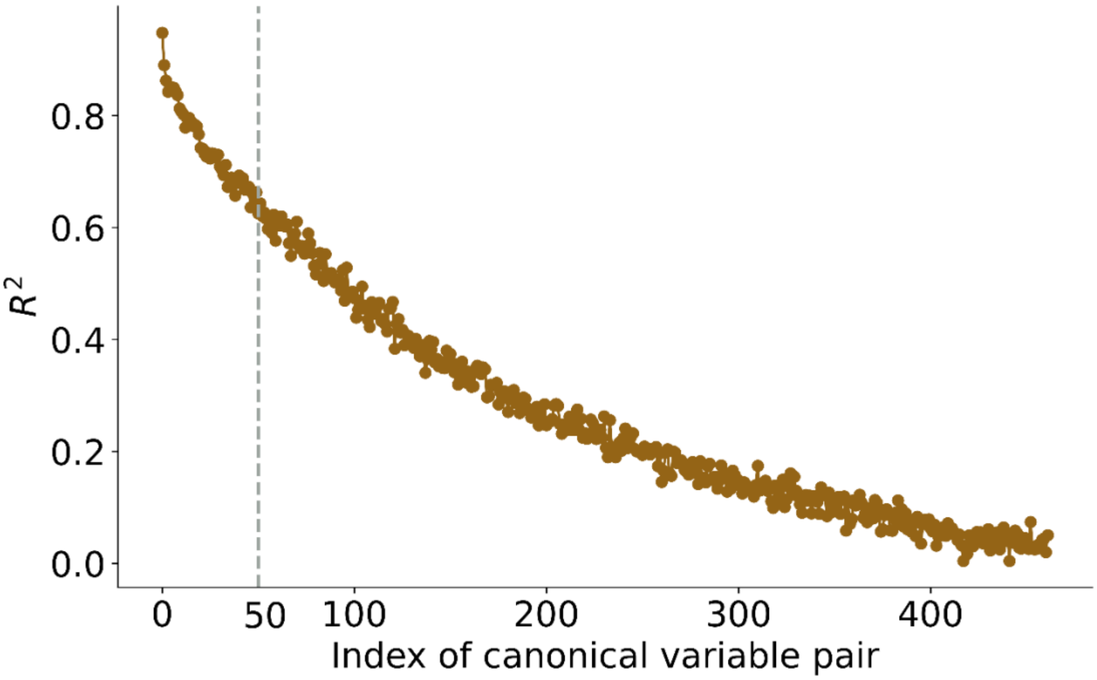


**Supplementary Figure 4.** The trend of the $R^2$ (squared Pearson correlation) between the Canonical Variable pairs of the decoder8 of GPT-2 and the encoder7 of Wav2Vec2.0—the most similar layer between these two models—varies with the index. We selected the top 50 canonical variables, whose $R^2$ values are around 0.6.
